# Polygenic Susceptibility of Aortic Aneurysms Associates to the Diameter of the Aneurysm Sac: the Aneurysm-Express Biobank Cohort

**DOI:** 10.1101/636795

**Authors:** Constance J.H.C.M. van Laarhoven, Jessica van Setten, Joost A. van Herwaarden, Dominique P.V. de Kleijn, Gerard Pasterkamp, Gert J. de Borst, Sander W. van der Laan

**Affiliations:** Department of Vascular Surgery, Division of Surgical Specialties, University Medical Center Utrecht, Utrecht University, Utrecht, the Netherlands; Laboratory of Clinical Chemistry and Hematology, Division Laboratories, Pharmacy, and Biomedical genetics, University Medical Center Utrecht, Utrecht University, Utrecht, the Netherlands; Cardiology, Division Heart & Lungs, University Medical Center Utrecht, Utrecht University, Utrecht, the Netherlands

**Author notes:** Correspondence and requests for reprint to:* Sander W. van der Laan PhD, Laboratory of Clinical Chemistry and Hematology, Division Laboratories, Pharmacy, and Biomedical Genetics, University Medical Center Utrecht, Utrecht University, room Fac03.02, PO Box 85500, 3508 GA Utrecht, the Netherlands. Ethical approval and informed consent*: All study patients were included from our Aneurysm-Express Biobank. The biobank has been approved by the local ethics committee, and all patients gave informed consent.

**Keywords:** Genetic risk variants, arterial aneurysm, polygenic risk score

## Abstract

**Purpose:** Abdominal aortic aneurysms (AAA) have a multifactorial pathology with both genetic and environmental risk factors. Recent genome-wide association studies (GWAS) have discovered ten genetic risk loci for AAA. To what extent these genetic loci contribute to the aneurysm pathology is yet unknown. This study aims to investigate whether genetic risk variants are associated with three clinical features: diameter of aneurysm sac, type of artery and symptoms.

**Methods:** We used aneurysm tissue from 415 patients included within the Aneurysm-Express biobank. A best fit polygenic risk score (PRS) based on previous GWAS effect size estimates was modeled for each clinical parameter by comparing model predictions across different *p*-value thresholds. Next, the established 10 risk variants for AAA were tested individually for association with selected clinical phenotypes. Models were corrected for age, sex, ancestral background, smoking status and diameter of the aneurysm sac or artery type if applicable, and data was normalized.

**Results:** The best fit PRS (including 272 SNPs with *P*_*T*_=0.01015) showed a significant correlation with diameter of the aneurysm sac (R^2^ = 0.019, *p* = 0.001). No association was found with clinical symptoms or type of artery. Individual variant analysis showed no clear associations with any of the clinical features.

**Conclusions:** Within the Aneurysm-Express Biobank Study, a weighted polygenic score of AAA susceptibility explained 1.9% of the phenotypic variation (*p* = 0.001) in aneurysm diameter. Individual risk variant analysis showed no clear associations. Given our limited sample size, future biobank collaborations need to confirm a potential causal role of individual SNPs on the pathology of aneurysms.

## Introduction

Abdominal aortic aneurysm disease (AAA) is a vascular pathology, affecting in particular elderly Western men. [1-3] Besides gender and smoking, a positive family history is a known predisposing factor for occurrence of the disease, [4-5] indicative of a strong heritable component to AAA.

In the past decade, genome-wide association studies (GWAS) have uncovered nine common genetic variants, *i.e.* single-nucleotide polymorphisms (SNPs), associated to AAA susceptibility. [6-12] While these GWAS point to part of the genetic underpinnings of AAA, the extent to which these variants influence the clinical presentation or how these loci contribute to pathology of the disease is still largely unknown.

Previous literature has shown many complex traits to be polygenic in origin, comprising small effects of hundreds or even thousands of common variants that in aggregate explain a substantial proportion of trait susceptibility and heritability. [13-15] For example, the International Schizophrenia Consortium [16] summarized weighted genetic effects [13] across nominally associated loci at increasingly liberal *p*-value thresholds into polygenic risk scores (PRS), and correlated PRS to disease susceptibility, demonstrating that a PRS can provide a reliable genetic indicator for clinical outcome.

In the present study, we investigated whether AAA susceptibility variants in aggregate are associated with three clinical phenotypes (maximum diameter of aneurysm sac, aneurysm-related symptoms, and type of artery) within the Aneurysm-Express Biobank Study [17] These phenotypes encompass intervention matters and are therefore clinically relevant. First, we constructed a weighted PRS based on summary level GWAS data for AAA [12] using increasingly liberal *p*-value thresholds and modeled a best fit PRS. Secondly, we tested the genome-wide significant risk SNPs for association with the selected clinical phenotypes in our cohort of patients with clinically manifested disease. Our results show that higher PRS associates to larger aneurysm diameter, but no association was found with clinical symptoms or type of artery.

## Methods

### Aneurysm-Express Biobank Study

The Aneurysm-Express is a biobank study that contains aneurysm sac tissue from patients undergoing open surgical repair of arterial aneurysms. The study design has been published previously. [17] Briefly, the study was approved by local medical ethics committee, eligible patients were operated in two different Dutch hospitals and all participants gave informed consent. All research was conducted according to the principles of the Declaration of Helsinki and its later amendments. For the present study we used clinical information from consecutive patients who were included between 2003 and 2013. The indications to perform open repair were based on at that time current guidelines. [3] Patients with arterial aneurysms caused by dissection, connective tissue disorders, and mycotic aneurysms or re-operated patients were excluded from this study. Risk factors and demographic data were obtained from clinical records and questionnaires at time of recruitment.

### Genotyping, quality control, and imputation

DNA of 503 patients in Aneurysm-Express Biobank Study was extracted from whole blood EDTA or (when no blood was available) aneurysm tissue samples following standardized in-house validated protocols for the Aneurysm-Express Genomics Study (AAAGS). Samples were sent for genotyping at the Genomic Analysis Center of the Helmholtz Zentrum Münich (Germany) according to OECD standards under study number M00750 using the Illumina HumanCoreExome BeadArray v1.1 (Illumina Inc., www.illumina.com).

Genotype calling was done with the GenomeStudio V2011.1 software and the Genotyping module version 1.9.4 using the original Illumina cluster and manifest files (humancoreexome-12v1-1_a.egt and HumanCoreExome-12-v1-1-C.bpm). The GenCall score cutoff was 0.15 as recommended by Illumina. The average call rate of all samples was 99.55% across 542,585 variants.

Subsequently, community standard quality control (QC) procedures were applied [18] to obtain high quality data. Samples with low average genotype calling and sex discrepancies (compared to the clinical data available) based on GenomeStudio metrics were excluded. The data was filtered on 1) individual (sample) call rate >97%, 2) SNP call rate >96%, 3) average heterozygosity rate ± 2.5 s.d., 4) relatedness (pi-hat > 0.20), 5) Hardy-Weinberg Equilibrium (HWE p < 1.0×10^−6^), and 6) population stratification excluding non-Europeans (based on 1000G phase 3). [19] zCall [20] was used to call missing exome-variants after QC. After QC and resulted in 478 samples and 541,569 variants (call rate = 99.99%) remained and were used for imputation.

Autosomal missing genotypes were imputed based on phased integrated data from 1000 Genomes (phase 3, version 5) and Genome of the Netherlands v5 [21] using IMPUTE2 (v2.3.0) [22] after pre-phasing genotyped data with SHAPEIT2 (v2.644). [23]

### Polygenic Risk Score (PRS)

We used the PRSice software [13] for creating the best fit PRS per phenotype by comparing scores across a range of different *p*-value thresholds. In short, PRSice calculates weighted PRS based on the estimated effects reported [12,39]: variants are pruned based on the linkage disequilibrium (r^2^ < 0.1, clump, range 500 kb) [20] as observed in AAAGS for bins of increasingly liberal *p*-value thresholds (*p*_*T*_, see Supplemental Table 1) preferentially keeping variants with lower *p*-values as reported by the meta-analysis of GWAS [12] (similar to the --clump algorithm in PLINK [24]). Odds ratios were natural log transformed to betas (β), we only included variants with MAF > 0.05 and imputation quality info-score > 0.8. In order to verify the predictive value of the PRS of the AAA-GWAS, we additionally modeled a PRS of summary statistics of the attention deficit hyperactivity disorder (ADHD) GWAS unrelated to AAA, [39] and tested this additional polygenic score for association with the selected clinical parameters in our study cohort.

### Individual SNP analysis

We selected nine SNPs that were identified in a recent GWAS meta-analysis for AAA. [12] These SNPs were tested for association with the clinical features separately. For the SNP lookup we used GWASToolKit (https://github.com/swvanderlaan/GWASToolKit, doi: 10.5281/zenodo.997862), which is a collection of scripts to execute SNPTEST v2.5.3 [25] analyses. Given the limited sample size, we calculated the expected power for this analysis. [26] In case of a risk allele frequency of <20%, and estimated OR of 1.10, the resulting power is ±80% (Supplemental Figure 1).

### Primary endpoints

The primary outcomes or phenotypes were artery type, symptoms, and diameter of the aneurysm sac. Artery type was defined as either aortic, or peripheral (iliac, femoral, popliteal, and carotid). Symptom status was defined as asymptomatic, or any aneurysm-related symptoms like thromboembolic events, local pain or swelling, or rupture of the aneurysm sac. Maximum diameter was measured by experienced radiologists at time of inclusion, on the double oblique plane of computed tomography angiography.

### Statistical analysis

PRS and individual SNP analyses were performed using linear and logistic regression models where appropriate, adjusted for sex, age, ancestral background using four principal components, smoking status, and diameter of the aneurysm sac or artery type if applicable. Nagelkerke’s r2 was used as a metric of the variance explained by the polygenic model. Statistical analyses were conducted using SPSS v25.0 (IBM Corp. Released 2017. IBM SPSS Statistics for Windows, Version 25.0. Armonk, NY: IBM Corp.), PRSice v2.1.4. [13], R v3.4.0 (R Core Team (2017), R: A language and environment for statistical computing. R Foundation for Statistical Computing, Vienna, Austria. URL http://www.R-project.org), and RStudio v3.4.1 (RStudio Team (2016). RStudio: Integrated Development for R. RStudio, Inc., Boston, MA URL www.rstudio.com).

## Results

In the present study, the Aneurysm-Express biobank comprised a total of 415 patients, average age of 69±8.1 years-old, whose majority (85%) is male, and 349 (84%) patients were treated for an AAA (Table 1).

**Table 1.** Aneurysm Express Biobank. Baseline characteristics.

| **Table 1. Aneurysm Express Biobank** |  |  |  |  |  |  |  |  |  |  |
| --- | --- | --- | --- | --- | --- | --- | --- | --- | --- | --- |
| Baseline characteristics. |  |  |  |  |  |  |  |  |  |  |
|  | **AAA**<br>**n = 349** |  | **Iliac**<br>**n = 13** |  | **Femoral**<br>**n = 9** |  | **Popliteal**<br>**n = 35** |  | **Carotid**<br>**n = 9** |  |
| **Male** | 292 | 84% | 13 | 100% | 8 | 100% | 34 | 97% | 4 | 44% |
| **Age at surgery in years (range)** | 70 | 7.1<br>(48-89) | 65 | 11.0<br>(48-86) | 67 | 11.2<br>(49-87) | 66 | 10.0<br>(45-83) | 62 | 9.7<br>(46-75) |
| **Aneurysm shape** |  |  |  |  |  |  |  |  |  |  |
| saccular | 16 | 5% | 0 | - | 0 | - | 1 | 3% | 3 | 33% |
| fusiform | 332 | 95% | 13 | 100% | 9 | 100% | 34 | 97% | 6 | 67% |
| **Reported aneurysm diameter (mm)** | 64 | 13.9<br>(31-118) | 46 | 12.4<br>(25-70) | 44 | 25.5<br>(24-100) | 35 | 20.0<br>(11-105) | 23 | 10.2<br>(12-38) |
| **Symptoms** |  |  |  |  |  |  |  |  |  |  |
| Ruptured | 28 | 8% | 1 | 8% | 0 | - | 2 | 6% | 0 | - |
| Any aneurysm related symptom | 82 | 24% | 4 | 31% | 4 | 44% | 19 | 54% | 6 | 68% |
| Asymptomatic | 237 | 68% | 7 | 54% | 5 | 56% | 14 | 40% | 3 | 33% |
| **Hypertension** | 275 | 79% | 7 | 54% | 4 | 44% | 22 | 63% | 4 | 44% |
| **Diabetes** | 58 | 17% | 1 | 8% | 1 | 11% | 4 | 11% | 1 | 11% |
| **Statin use** | 237 | 68% | 7 | 54% | 3 | 33% | 19 | 54% | 6 | 67% |
| **BMI** | 25.9 | 4.1 | 26.4 | 5.2 | 27.7 | 4.6 | 27.2 | 3.6 | 23.7 | 2.6 |
| **Smoking** | 126 | 36% | 6 | 46% | 2 | 22% | 12 | 34% | 0 | - |
| Data are given as numbers (percentage) or mean (standard deviation). |  |  |  |  |  |  |  |  |  |  |
| Abbreviations: AAA = abdominal aortic aneurysm, mm = millimeter, BMI = body mass index. |  |  |  |  |  |  |  |  |  |  |

We used summary statistics from the largest meta-GWAS for AAA [12] so far to calculate weighted polygenic scores at increasingly liberal *p*-value thresholds (*p*_*T*_, Supplemental Table 1) and correlated these to artery type, symptoms, and diameter of the aneurysm sac (Figure 1A-C). A PRS including 272 variants at *p*_*T*_ of 0.01015 explained the largest proportion in diameter (R^2^ = 0.019, *p* =0.001, Supplemental Table 2). Distributions of the AAA-PRS for diameter was comparable between artery types (Kruskal-Wallis test *p*-value = 0.135, Supplemental Figure 2). No association was found for artery type and symptom status (Supplemental Table 2). We verified all AAA-PRS findings with another unrelated PRS derived from a ADHD GWAS, [39] and correlated these with the selected clinical parameters in our study cohort (Figure 1D-F). No associations with the ADHD-PRS were observed (Figure 1D-F, Supplementary Table 3). Next, we tested ten known AAA SNPs for association with diameter of the aneurysm sac (Table 2), artery type and symptom status (Supplemental Table 3). For diameter, a nominal association was found with rs1466535 (12q13.3, *LRP1, p* = 0.013). For rs602633 (1p13.3, *PSRC1-CELSR2-SORT1*), rs1795061 (1q32.3, *SMYD2*), rs2836411 (21q22.2, *ERG*) and rs1466535 a concordant effect direction was observed. For six risk variants, the effect direction was disconcordant with the GWAS results.

**Figure 1.**
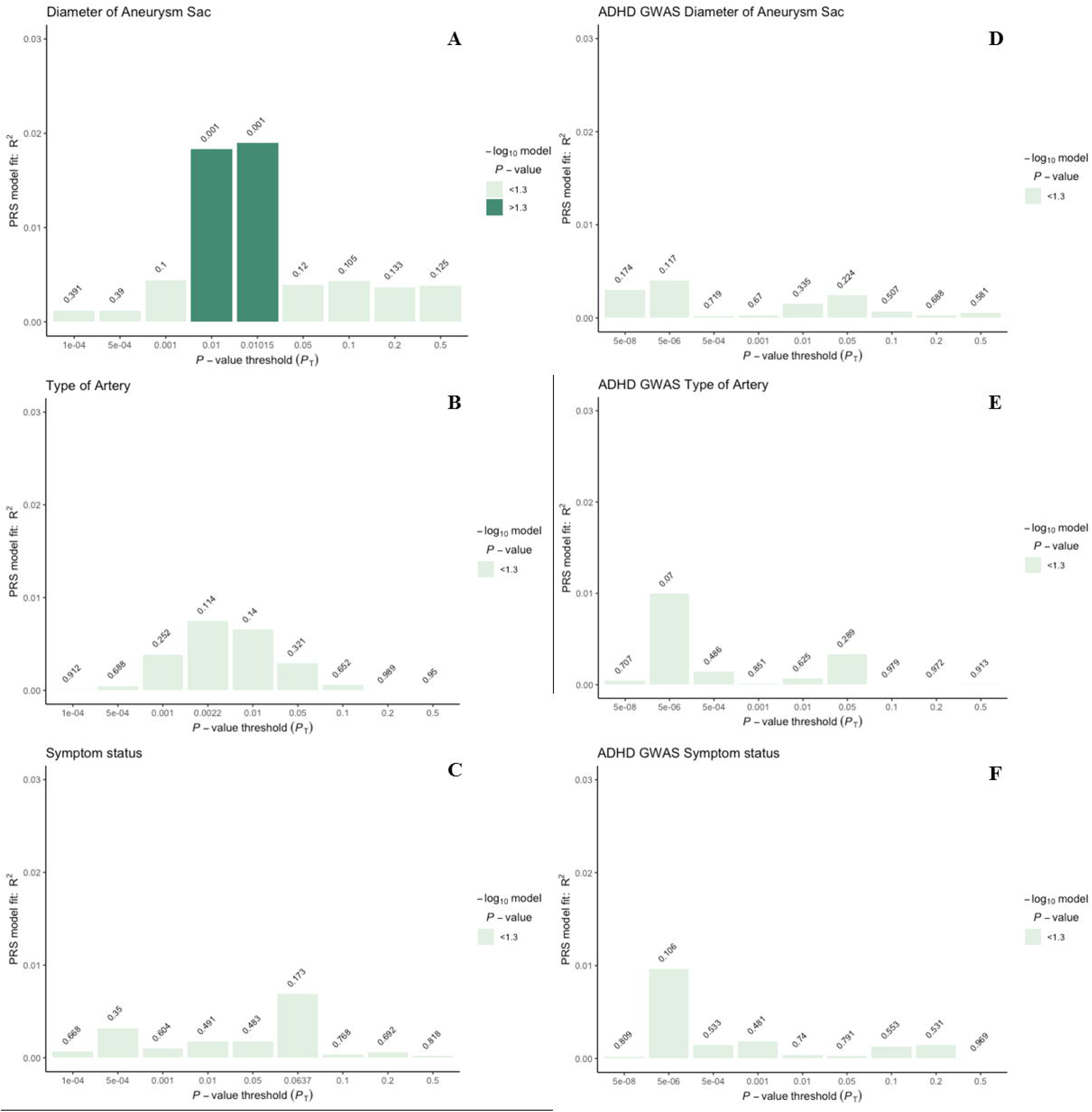
PRSice [14] generated weighted model for: **A** Diameter of the aneurysm sac, **B** Type of artery, and **C** Symptom status. Models are summarized in Supplementary Table 2&3. **D-E-F** indicate unrelated PRS derived from ADHD GWAS summary statistics [39], showing no association with any of the selected clinical phenotypes in the Aneurysm-Express Biobank cohort. *Abbreviations*: PRS = polygenic risk score, ADHD = attention deficit hyperactivity disorder, GWAS = genome-wide association study.

**Table 2.**
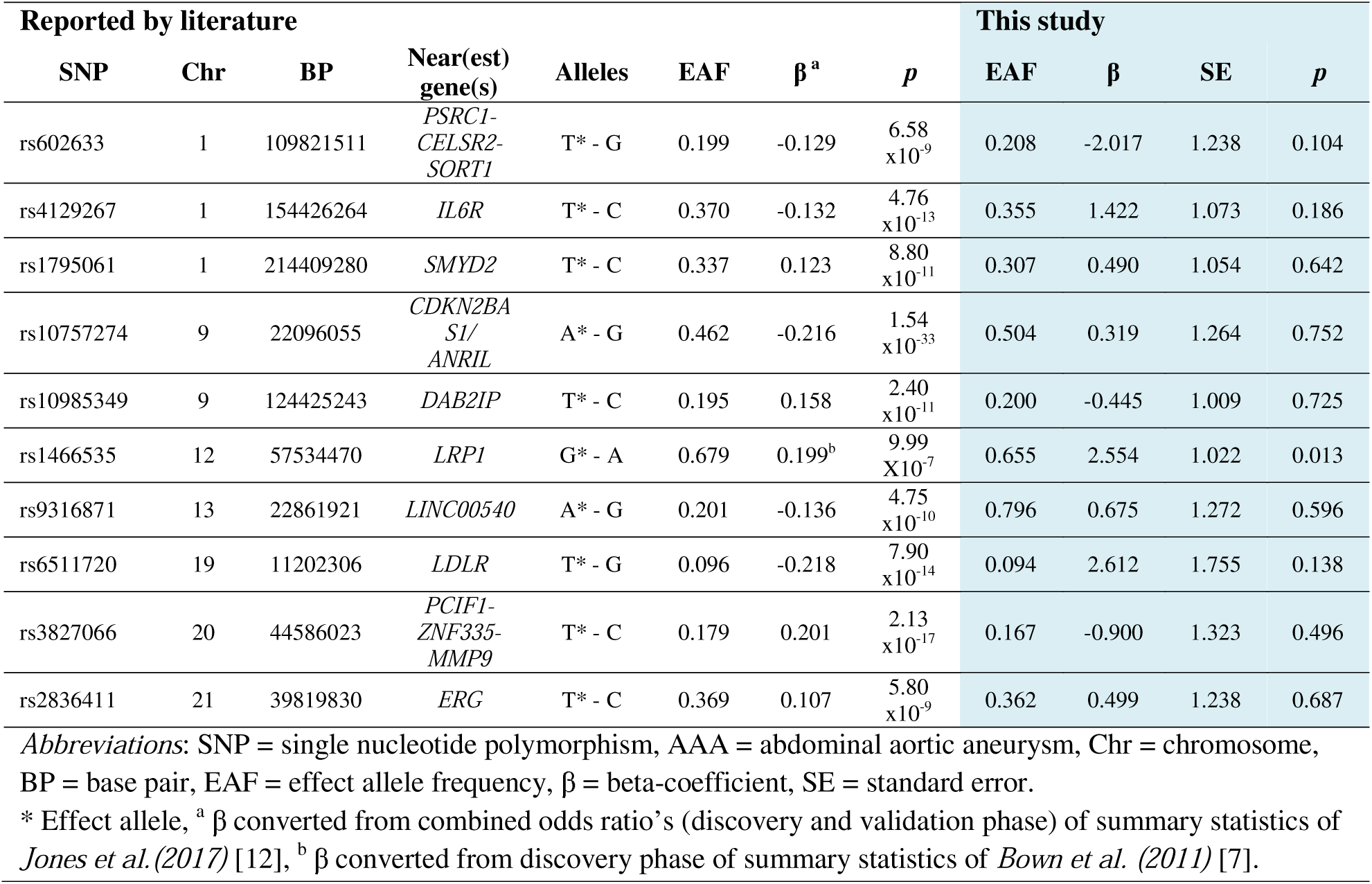
Individual SNP analysis. AAA associated SNPs reported by GWAS and the association results for diameter of the aneurysm sac (in millimeters).

## Discussion

In this study, we investigated the association of genetic susceptibility variants with clinical phenotypes within a biobank consisting of both aortic and peripheral aneurysms. We demonstrated that within our Aneurysm-Express biobank cohort, polygenic scores of AAA susceptibility was associated with diameter of the aneurysm sac. No association was found with clinical symptoms or type of artery.

The clinical translation of genetic research has been a topic of interest for the last decades. For rare diseases like Marfan or Ehlers-Danlos syndrome, genetic clinical utility has been proven and applied, [27] but for more common complex diseases like AAA, the interpretation and translation of large-scale genetic studies is challenging at best. [28] GWAS-derived association studies were mainly investigated in intracranial and abdominal aneurysm patients. [29-34] The majority of these studies used a genetic risk score (GRS) based on the index SNPs from GWAS to study additional genetic predicting value to models consisting of demographic characteristics and health parameters. A previous study using a GRS including 4 variants showed that a high GRS was associated with aneurysm growth rate independent of baseline abdominal aortic size. [38] Recently, a similarly calculated, but polygenic risk approach has proved clinical utility in atherosclerosis, statin therapy, and breast cancer. [35-37] A polygenic approach can optimize power by using a liberal *p*-value threshold, enforced by the evidence that additive weakly effects of nominally associated common variants explain part of the heritability of, and susceptibility to common complex diseases. [14,15] Our study uses a polygenic risk approach within a cohort comprising both aortic and peripheral aneurysm patients and shows that a high polygenic risk of AAA is associated to larger diameter of the aneurysm sac.

After correcting for multiple testing, no relevant effects of individual SNPs were observed, and the majority of the effect directions in our analysis are discordant with the recent GWAS meta-analysis. [12] This is probably mainly due to limited size and power of both the published GWAS (4,972 cases and 99,858 controls) and our present study (Supp. Figure 2). Differences could also be explained by differences in inclusion criteria; the GWAS analysis is performed in patients with AAA ≥ 30mm, including smaller AAA that have been followed-up and was compared to healthy subjects. [28] In contrast, our cohort consists of only patients including symptomatic patients or progressive aneurysmal disease that requires surgical treatment. For this, it is arguable that other risk variants have a more prominent role in further deterioration of the vessel wall and have another direction than GWAS identified loci that may contribute particularly to initiation of aneurysmal disease.

Whether aortic and intracranial aneurysms share genetic susceptibility, was earlier investigated in a GWAS of four cohorts. [31] These studies provide evidence that variants at 9p21, 18q11, 15q21, and 2q33 are consistently associated with intracranial, thoracic aneurysms and AAA. Their analysis revealed no additional risk loci associated with joint aneurysms, presumably by their limited sample size of 3094 cases and 9521 controls. The PRS distribution of diameter was comparable across artery types (Supplemental Figure 2), this suggests overlapping genetic pathways between peripheral aneurysms and central aneurysms.

The Aneurysm-Express biobank cohort is limited in size when compared to other biobanks, yet unique in its scope, given the availability of arterial aneurysm samples from the open arterial surgery era for analyses. Future studies should focus on biobank collaborations, enrichment, and harmonization to facilitate the in-depth scrutiny and replication of genetic susceptibility on AAA pathology. Our biobank is surgically driven, and measured diameters may cluster around surgical intervention thresholds (e.g. for AAA ≥ 55 mm) limiting the generalizability, still we observed normally distributed diameters.

## Conclusions

In conclusion, we show that a weighted polygenic score of AAA susceptibility explained 1.9% of the phenotypic variation (*p* = 0.001) in aneurysm diameter in the Aneurysm-Express Biobank Study. Future studies should focus on biobank collaborations, enrichment, and harmonization to assess potential impact of known susceptibility loci on the pathology of AAA.

