## Supplemental material for "Polygenic Susceptibility of Aortic Aneurysms Associates to the Diameter of the Aneurysm Sac: the Aneurysm-Express Biobank Cohort"

#### Appendix:

- Supplemental Figure 1
- Supplemental Figure 2
- Supplemental Table 1
- Supplemental Table 2
- Supplemental Table 3
- Supplemental Table 4

### Supplemental Figure 1

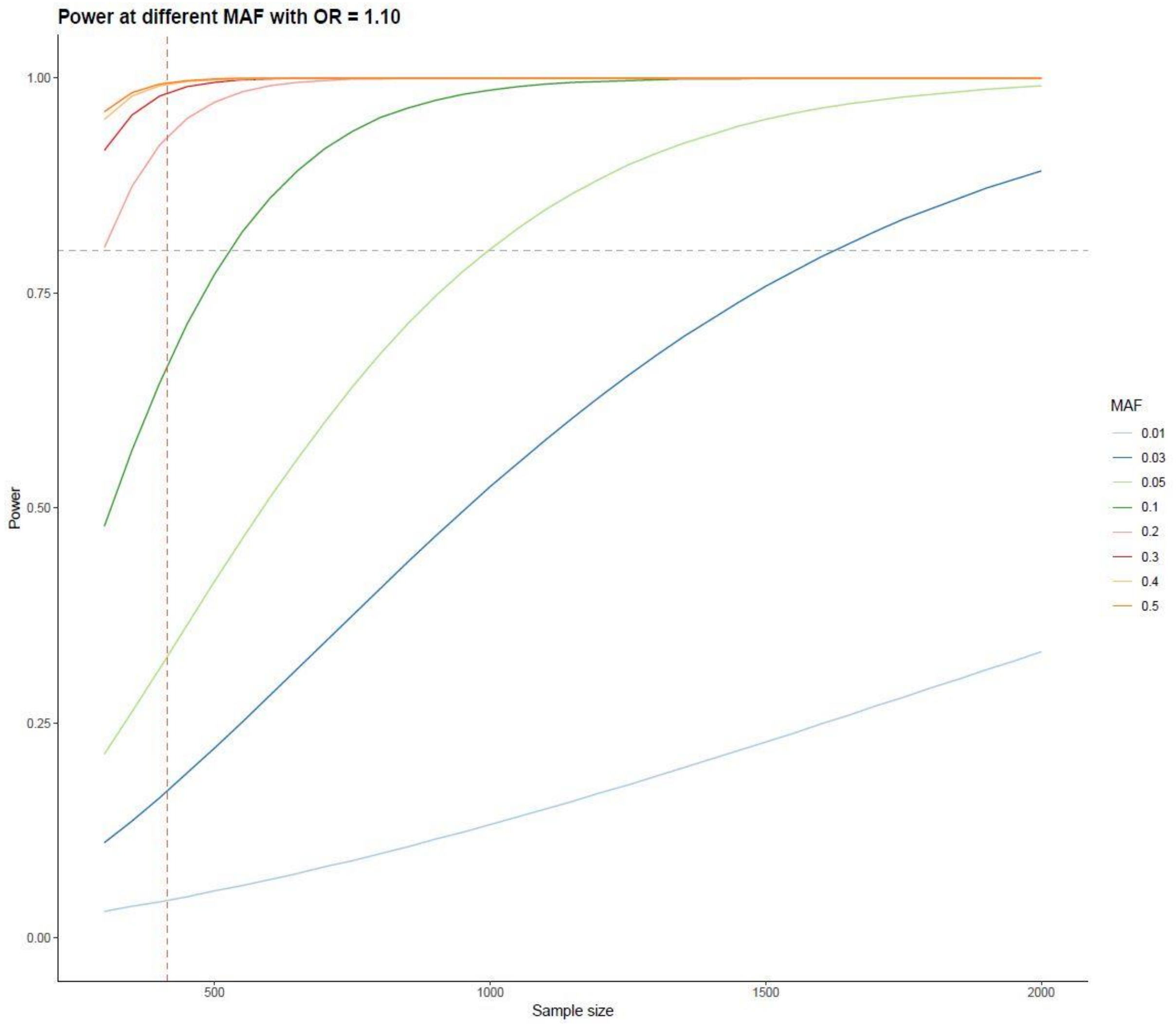


**Supplemental Figure 1.** Plot indicating study power estimates at different minor allele frequencies (MAF), with an average odds ratio (OR) of 1.10. Each solid colored line indicates a different MAF. Red dashed line indicates present study size of 415 samples, grey dashed line indicates a power estimate of ±80% for risk variants MAF < 0.2.

#### Supplemental Figure 2

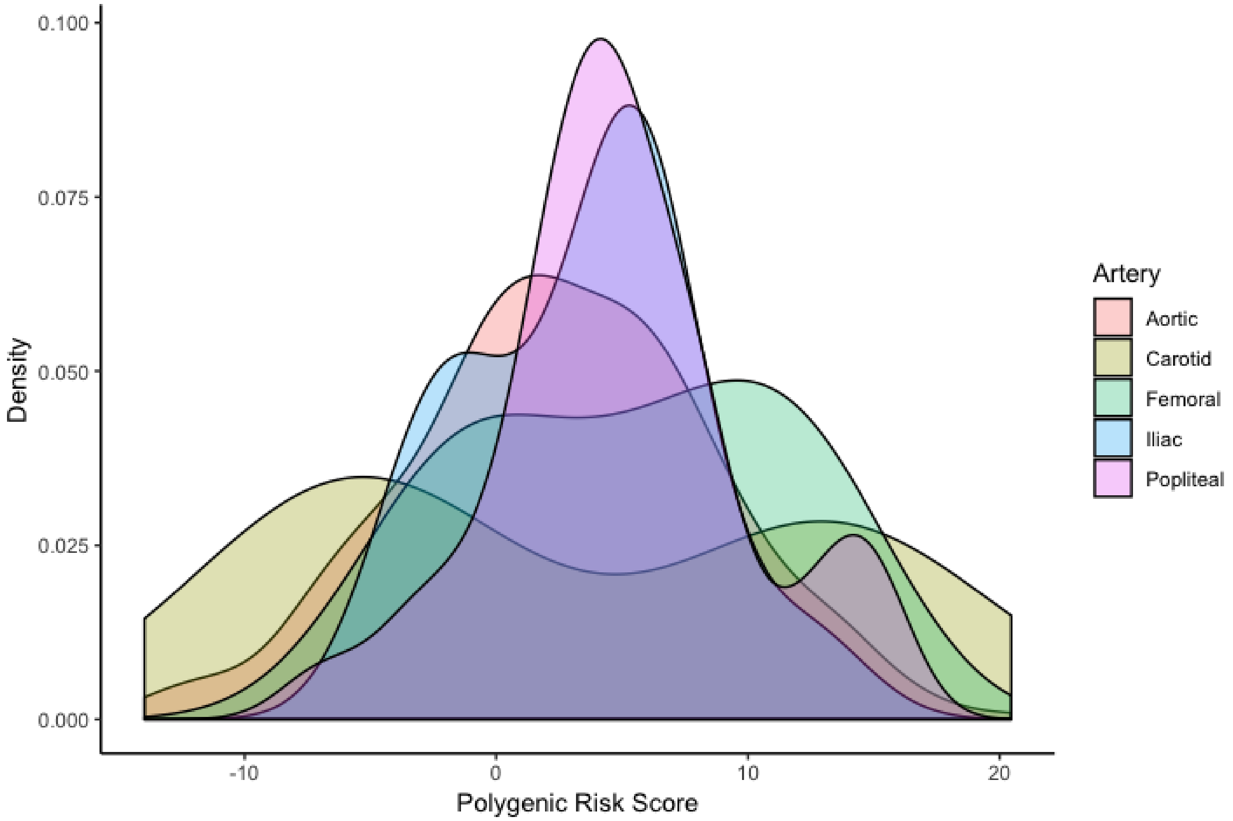


**Supplemental Figure 2.** Overlay density plot of polygenic burden/risk score (PRS) for diameter per artery type. Distribution of PRS is comparable per artery type (*Kruskal-Wallis test p*-value = 0.135).

#### Supplemental Table 1. Polygenic models.

A baseline model was constructed correcting for age, sex, ancestral background using four principal components, smoking status, and maximum diameter of the aneurysm sac or artery type if applicable. $p_T$: the GWAS p-value threshold of association.

| **Model** | **Includes** |
|---|---|
| GWAS | Baseline + GWAS hits $p_T < 5\text{x}10^{-8}$ |
| $p < 5\text{x}10^{-6}$ | Baseline + all SNPs $p_T < 5\text{x}10^{-6}$ |
| $p < 5\text{x}10^{-4}$ | Baseline + all SNPs $p_T < 5\text{x}10^{-4}$ |
| $p < 0.001$ | Baseline + all SNPs $p_T < 0.001$ |
| $p < 0.01$ | Baseline + all SNPs $p_T < 0.01$ |
| $p < 0.05$ | Baseline + all SNPs $p_T < 0.05$ |
| $p < 0.1$ | Baseline + all SNPs $p_T < 0.1$ |
| $p < 0.2$ | Baseline + all SNPs $p_T < 0.2$ |
| $p < 0.5$ | Baseline + all SNPs $p_T < 0.5$ |

**Supplemental Table 2. Detailed polygenic risk score (PRS) models derived from AAA GWAS [12] for selected clinical phenotypes.**

**Diameter of the aneurysm sac**

| $p_T$ | $R^2$ | β | SE | *p*-value | Number of SNPs |
|---|---|---|---|---|---|
| $5x10^{-8}$ | 0.001 | -1.932 | 2.143 | 0.368 | 1 |
| $5x10^{-6}$ | 0.000 | 0.075 | 1.419 | 0.958 | 3 |
| $5x10^{-4}$ | 0.001 | 0.371 | 0.431 | 0.390 | 24 |
| 0.001 | 0.004 | 0.493 | 0.299 | 0.100 | 40 |
| 0.01 | 0.018 | 0.394 | 0.116 | 0.001* | 267 |
| **0.01015** | **0.019** | **0.398** | **0.115** | **0.001*** | **272** |
| 0.05 | 0.004 | 0.095 | 0.061 | 0.120 | 1,008 |
| 0.1 | 0.004 | 0.075 | 0.046 | 0.105 | 1,634 |
| 0.2 | 0.004 | 0.054 | 0.036 | 0.133 | 2,494 |
| 0.5 | 0.004 | 0.045 | 0.029 | 0.125 | 4,086 |

**Artery type**

| $p_T$ | $R^2$ | β | SE | *p*-value | Number of SNPs |
|---|---|---|---|---|---|
| $5x10^{-8}$ | 0.005 | -0.886 | 0.681 | 0.193 | 1 |
| $5x10^{-6}$ | 0.003 | -0.382 | 0.411 | 0.353 | 3 |
| $5x10^{-4}$ | 0.000 | 0.055 | 0.138 | 0.688 | 24 |
| 0.001 | 0.004 | 0.110 | 0.096 | 0.252 | 39 |
| **0.0022** | **0.007** | **0.099** | **0.063** | **0.114** | **88** |
| 0.01 | 0.007 | 0.056 | 0.038 | 0.140 | 267 |
| 0.05 | 0.030 | 0.020 | 0.020 | 0.321 | 1,008 |
| 0.1 | 0.001 | 0.006 | 0.014 | 0.652 | 1,632 |
| 0.2 | 0.000 | 0.000 | 0.012 | 0.989 | 2,493 |
| 0.5 | 0.000 | -0.001 | 0.009 | 0.950 | 4,086 |

**Symptom status**

| $p_T$ | $R^2$ | β | SE | *p*-value | Number of SNPs |
|---|---|---|---|---|---|
| $5x10^{-8}$ | 0.000 | 0.032 | 0.342 | 0.925 | 1 |
| $5x10^{-6}$ | 0.000 | 0.023 | 0.226 | 0.921 | 3 |
| $5x10^{-4}$ | 0.003 | 0.064 | 0.069 | 0.350 | 24 |
| 0.001 | 0.001 | 0.025 | 0.049 | 0.604 | 39 |
| 0.01 | 0.002 | 0.013 | 0.019 | 0.491 | 270 |
| 0.05 | 0.002 | 0.007 | 0.010 | 0.483 | 1,009 |
| **0.0637** | **0.007** | **0.012** | **0.009** | **0.173** | **1,213** |
| 0.1 | 0.000 | 0.002 | 0.007 | 0.768 | 1,632 |
| 0.2 | 0.001 | 0.002 | 0.006 | 0.692 | 2,494 |
| 0.5 | 0.000 | 0.001 | 0.005 | 0.818 | 4,086 |

**Bold** indicates best fitted model, * indicates $p < 0.05$.

*Abbrevations:* PRS = polygenic risk score. $p_T$ = *p*-value threshold. β = beta coefficient. SE = standard error, SNPs = single nucleotide polymorphisms.

#### Supplemental Table 3. Detailed polygenic risk score (PRS) models derived from attention deficit hyperactivity disorder GWAS [39] for selected clinical phenotypes.

**Diameter of the aneurysm sac**

| $p_T$ | $R^2$ | β | SE | *p*-value | Number of SNPs |
|---|---|---|---|---|---|
| $5x10^{-8}$ | 0.003 | -0.973 | 0.714 | 0.174 | 10 |
| $5x10^{-6}$ | 0.004 | -1.105 | 0.703 | 0.117 | 77 |
| $5x10^{-4}$ | 0.000 | -0.256 | 0.710 | 0.719 | 1,179 |
| 0.001 | 0.000 | 0.306 | 0.716 | 0.670 | 1,858 |
| 0.01 | 0.002 | -0.702 | 0.727 | 0.335 | 8,155 |
| 0.05 | 0.002 | -0.907 | 0.745 | 0.224 | 20,395 |
| 0.1 | 0.001 | -0.494 | 0.745 | 0.507 | 28,657 |
| 0.2 | 0.000 | -0.298 | 0.741 | 0.688 | 39,167 |
| 0.5 | 0.000 | -0.409 | 0.741 | 0.581 | 55,850 |

**Artery type**

| $p_T$ | $R^2$ | β | SE | *p*-value | Number of SNPs |
|---|---|---|---|---|---|
| $5x10^{-8}$ | 0.000 | 0.086 | 0.230 | 0.707 | 10 |
| $5x10^{-6}$ | 0.010 | -0.424 | 0.234 | 0.070 | 77 |
| $5x10^{-4}$ | 0.001 | -0.168 | 0.240 | 0.486 | 1,179 |
| 0.001 | 0.000 | -0.045 | 0.239 | 0.851 | 1,858 |
| 0.01 | 0.001 | -0.115 | 0.235 | 0.625 | 8,155 |
| 0.05 | 0.003 | -0.254 | 0.239 | 0.289 | 20,395 |
| 0.1 | 0.000 | -0.006 | 0.226 | 0.979 | 28,657 |
| 0.2 | 0.000 | -0.008 | 0.232 | 0.972 | 39,167 |
| 0.5 | 0.000 | -0.026 | 0.234 | 0.913 | 55,850 |

**Symptom status**

| $p_T$ | $R^2$ | β | SE | *p*-value | Number of SNPs |
|---|---|---|---|---|---|
| $5x10^{-8}$ | 0.000 | 0.028 | 0.115 | 0.809 | 10 |
| $5x10^{-6}$ | 0.010 | -0.184 | 0.114 | 0.106 | 77 |
| $5x10^{-4}$ | 0.001 | -0.070 | 0.113 | 0.533 | 1,179 |
| 0.001 | 0.002 | -0.081 | 0.114 | 0.481 | 1,858 |
| 0.01 | 0.000 | -0.038 | 0.116 | 0.740 | 8,155 |
| 0.05 | 0.000 | -0.032 | 0.119 | 0.791 | 20,395 |
| 0.1 | 0.001 | -0.071 | 0.120 | 0.553 | 28,657 |
| 0.2 | 0.001 | -0.074 | 0.119 | 0.531 | 39,167 |
| 0.5 | 0.000 | -0.005 | 0.118 | 0.969 | 55,850 |

**Bold** indicates best fitted model, * indicates $p < 0.05$.
*Abbrevations:* $p_T$ = *p*-value threshold. β = beta coefficient. SE = standard error, SNPs = single nucleotide polymorphisms.

### Supplemental Table 4

**Supplemental Table 4.**

AAA associated SNPs reported by GWAS and the association results for artery type and symptom status.

| **Reported by literature** | | | | | | | | **This study** | | | | | | | |
|---|---|---|---|---|---|---|---|---|---|---|---|---|---|---|---|
| | | | | | | | | *Artery type* | | | | *Symptom status* | | | |
| **SNP** | **Chr** | **BP** | **Near(est) gene(s)** | **Alleles** | **EAF** | **β [a]** | ***p*** | **EAF** | **β** | **SE** | ***p*** | **EAF** | **β** | **SE** | ***p*** |
| rs602633 | 1 | 109821511 | *PSRC1-CELSR2-SORT1* | T* - G | 0.199 | -0.129 | $6.58 \times 10^{-9}$ | 0.208 | 0.034 | 0.306 | 0.911 | 0.208 | -0.694 | 0.192 | 0.132 |
| rs4129267 | 1 | 154426264 | *IL6R* | T* - C | 0.370 | -0.132 | $4.76 \times 10^{-13}$ | 0.355 | 0.173 | 0.270 | 0.522 | 0.361 | 0.564 | 0.171 | 0.133 |
| rs1795061 | 1 | 214409280 | *SMYD2* | T* - C | 0.337 | 0.123 | $8.80 \times 10^{-11}$ | 0.307 | 0.149 | 0.276 | 0.591 | 0.315 | -0.550 | 0.165 | 0.125 |
| rs10757274 | 9 | 22096055 | *CDKN2BAS 1/ANRIL* | A* - G | 0.462 | -0.216 | $1.54 \times 10^{-33}$ | 0.504 | -0.015 | 0.263 | 0.955 | 0.506 | -0.377 | 0.160 | 0.276 |
| rs10985349 | 9 | 124425243 | *DAB2IP* | T* - C | 0.195 | 0.158 | $2.40 \times 10^{-11}$ | 0.200 | -0.015 | 0.339 | 0.964 | 0.204 | 0.149 | 0.200 | 0.711 |
| rs1466535 | 12 | 57534470 | *LRP1* | G* - A | 0.679 | 0.199[b] | $9.99 \times 10^{-7}$ | 0.655 | 0.039 | 0.267 | 0.885 | 0.655 | -0.255 | 0.166 | 0.124 |
| rs9316871 | 13 | 22861921 | *LINC00540* | A* - G | 0.201 | -0.136 | $4.75 \times 10^{-10}$ | 0.796 | 0.176 | 0.348 | 0.613 | 0.794 | 0.037 | 0.197 | 0.930 |
| rs6511720 | 19 | 11202306 | *LDLR* | T* - G | 0.096 | -0.218 | $7.90 \times 10^{-14}$ | 0.094 | 0.306 | 0.469 | 0.514 | 0.098 | 0.069 | 0.273 | 0.902 |
| rs3827066 | 20 | 44586023 | *PCIF1-ZNF335-MMP9* | T* - C | 0.179 | 0.201 | $2.13 \times 10^{-17}$ | 0.167 | 0.190 | 0.345 | 0.583 | 0.176 | -0.879 | 0.207 | 0.048 |
| rs2836411 | 21 | 39819830 | *ERG* | T* - C | 0.369 | 0.107 | $5.80 \times 10^{-9}$ | 0.361 | 0.433 | 0.311 | 0.164 | 0.364 | -0.500 | 0.194 | 0.224 |

*Abbreviations*:AAA = abdominal aortic aneurysm, SNP = single nucleotide polymorphism, Chr = chromosome, BP = base pair, EAF = effect allele frequency, β = beta-coefficient, SE = standard error.

* Effect allele, [a] β converted from combined odds ratio's (discovery and validation phase) of summary statistics of *Jones et al.(2017)* [12].

[b] β converted from discovery phase of summary statistics of *Bown et al. (2011)* [7].
